# Statistical optimization of culture conditions to improve cell density of *Komagataeibacter medellinensis* NBRC 3288: the first step towards to optimize bacterial cellulose production

**DOI:** 10.1101/381186

**Authors:** Carlos Molina-Ramírez, Robin Zuluaga, Cristina Castro, Piedad Gañán

## Abstract

The cell density yield of *Komagataeibacter medellinensis* NBRC 3288 was optimized applying response surface methodology (RSM) for the first time with this strain. Four factors were evaluated: glucose, initial pH, MgSO_4_ and KH_2_PO_4_, these were analyzed in a fractional factorial design to verify which were important to cell density yield. The best treatment assayed showed a yield of 1.09 OD_600_ and analysis of variance (ANOVA) selected glucose, inital pH and KH_2_PO_4_ as the significant factors. Next, a new experimental region with center point at glucose 0.5 %(wt./v), initial pH at 5.58 and KH_2_PO_4_ 0.058 %(wt./v) was obtained by steepest ascent method and used to optimize biomass yield by Box-Behnken design. The results indicate that the optimum culture medium conditions predicted by the mathematical model (R^2^_adj_=72.09 %) were: glucose, 0.54 %(wt./v); peptone, 0.5 %(wt./v); yeast extract, 0.5 %(wt./v); KH_2_PO_4_, 0.059 %(wt./v); MgSO_4_, 0.025 %(wt./v); NaH_2_PO_4_, 0.267 %(wt./v) and pH, 5.18 (adjusted with citric acid, 0.2 %(wt./v)) to obtain an optimum cell density yield of 2.85 OD_600_ with an improve of 68 % respect to the best result obtained at the begin, as well as μ_m_ improved 34.78 % and a reduction in cost of culture medium due to glucose concentration reduction of 75 %.

**Importance:** Despite the important role of producer-strain play in bacterial cellulose production, until now there is not anu report about how to improve cell density, as a primary stage, that allows reach higher production of BC in later stages. In this study, was optimized the growth conditions of *K. medellinensis* (a BC producer in acid pH) in order to increase the cell density, as the primer stage in the whole process to optimize the production of BC, which is a very important supplie in different fields of science such as: food, materials, water treatment, tissue engineering, etc. The optimization process was achieved with savings in production costs of the culture medium, through the reduction of glucose concentration by 75 % and through the reduction of the microorganism’s growth time.

## Introduction

*Komagataeibacter medellinensis* NBRC 3288 belongs to a group of obligate aerobic bacteria named acetic acid bacteria (AAB), which are characterized for converting sugars and alcohols into their corresponding organic acids through of a incomplete oxidative fermentation (Deppenmeier and Ehrenreich 2009). However, *Komagataeibacter medellinensis* NBRC 3288 is characterized also for producing cellulose, namely bacterial cellulose (BC), at very low pH level (Castro et al. 2012).

BC is a polymer formed by units of glucose linkaged by β-1,4 glycosidic linkage with a huge potential in different industries such as food (Shi et al. 2014), paper (Santos et al. 2016), medical (Thomas et al. 2013) or composite materials (Shah et al. 2013). BC biogenesis is a four-step process that begins with phosphorylation of glucose and finish with cellulose synthesis (Haigler and Benziman 1982; Ross et al. 1991; Zhang 2013), this process of leading from the hexose phosphate substrate to sugar nucleotides, represent the interface between primary metabolism and secondary metabolism (Ramos et al. 2001) suggesting that BC is a secondary metabolite beacuse is not associated to energetic metabolism of *Komagataeibacter medellinensis* NBRC 3288. Initial cell density is a key factor to improve BC production as was demonstrated by Okiyama et al. (Okiyama et al. 1992) and Zeng et al. (Zeng et al. 2011) and reduce the time for this stage is a remarkable fact that could reduce costs of BC production process guaranteeing a high density of cells as a previous step of BC optimization.

Surface response methodology (RSM) is a versatile and powerful tool widely used in many field of investigation (Shi et al. 2006; Bezerra et al. 2008; Hajati et al. 2015). Despite this, a reference analysis suggest that this is the first study in optimizing a high-density cultivation conditions for the strain *K. medellinensis* NBRC 3288.

In this work the aim is improve cell density of *K. medellinensis* NBRC 3288 through response surface methodology (RSM) and offer a new method to obtain the inoculum in order to optimize BC production as the main goal.

## Materials and methods

### Microorganism and seed culture

*Komagataeibacter medellinensis* NBRC 3288 was obtained from homemade vinegar in Medellín, Colombia (Castro et al. 2013). The strain was stored in vials in HS medium (Hestrin and Schramm 1954) at pH 3.5 and glycerol 20 (\%v/v) at −80 °C. Seed culture medium contained %(wt./v): glucose, 2; peptone, 0.5; yeast extract, 0.5; Na_2_HPO_4_, 0.267; citric acid, 0.8 and Celluclast 1.5L at 3 %(v/v) added after sterilization.

To obtain seed culture, one vial was added to 100 mL of seed culture medium in agitated conditions on orbital shaker at 120 rpm for 2 days.

### Fermentation screening stage

After 2 days, seed culture was centrifuged at 9000 rpm for 20 min at 7-8 °C and pellet was collected and resuspended in sterilized distilled water until adjust the OD at 0.5 using a UV spectrophotometer at 600 nm OD_600_). Then an inoculum size of 10 %(v/v) were transferred into shake flask with 90 mL of culture media with different conditions at 28 °C for 120 h.

### Fermentation optimization stage

Fermentation was carried out at same conditions described above, only fermentation time was changed until 72 h. At this time start decreasing the biomass growth and will be the growth stage at which will take inoculum for bacterial cellulose production because its nature of secondary metabolite.

### Optimization by RSM

A fractional factorial design 2^(4–1)^ was performed as screening design to select the significant factors. Four factors were evaluated: glucose concentration, initial pH, KH_2_PO_4_ concentration and MgSO_4_ concentration. Once defined significant factors, was applied steepest ascent methodology to get the experimental optimum region more accurately. The levels of factors are shown in Table 1 and the response was OD_600_ at 72 h of fermentation.

**Table 1.**
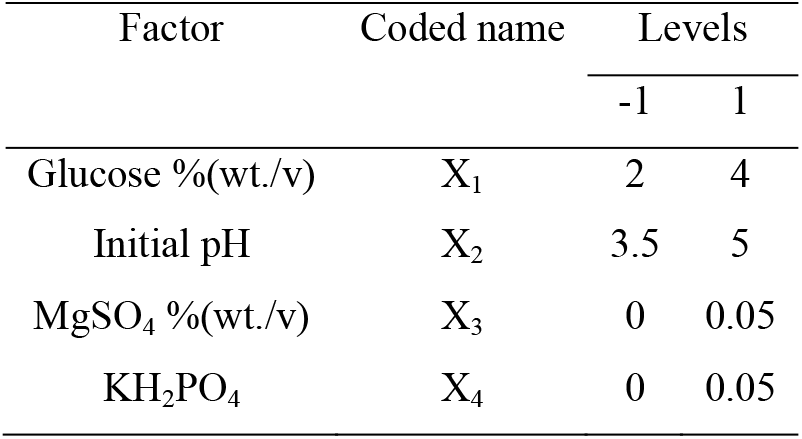
Levels factors evaluated in screening design

| Factor | Coded name | Levels |  |
| --- | --- | --- | --- |
|  |  | -1 | 1 |
| Glucose %(wt./v) | X<sub>1</sub> | 2 | 4 |
| Initial pH | X<sub>2</sub> | 3.5 | 5 |
| MgSO<sub>4</sub> %(wt./v) | X<sub>3</sub> | 0 | 0.05 |
| KH<sub>2</sub>PO<sub>4</sub> | X<sub>4</sub> | 0 | 0.05 |

Once the significant factors were selected in screening design and a new center point was selected by steepest ascent methodology, a Box-Behnken experimental design was applied to obtain a second order model that describe the experimental data, at least 70 % expressed by R^2^ parameter (Gutiérrez Pulido et al. 2004). The mathematical model obtained for three factors is expressed in the form:

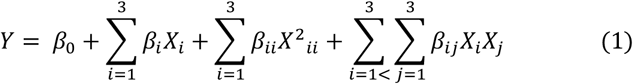

Where β_0_ is the intercept coefficient, β_i_ are the linear coefficients, β_ii_ are the quadratic coefficients and β_ij_ are the interactions coefficients. The statistical analysis was performed through analysis of variance (ANOVA).

The design of experiments, the statistical analysis and the optimization process were performed with the statistical software package STATGRAPHICS^®^ Centurion 64 XVI, StatPoint Technologies, Inc, The Plains, Virginia, USA.

## Results

### Significant factors selection

*K. medellinensis* NBRC 3288 growth behavior was evaluated by changing levels in 4 factors, namely glucose, initial pH, KH_2_PO_4_ and MgSO_4_. Figure 1 shows the effect of the 8 different experimental treatment on growth profile of *K. medellinensis* NBRC 3288 and the fitted line using modified Gompertz’s model (Zwietering et al. 1990). According to Figure 1, at 72 h begins deceleration stage and the results of biomass production are shown in Table 2.

**Figure 1.**
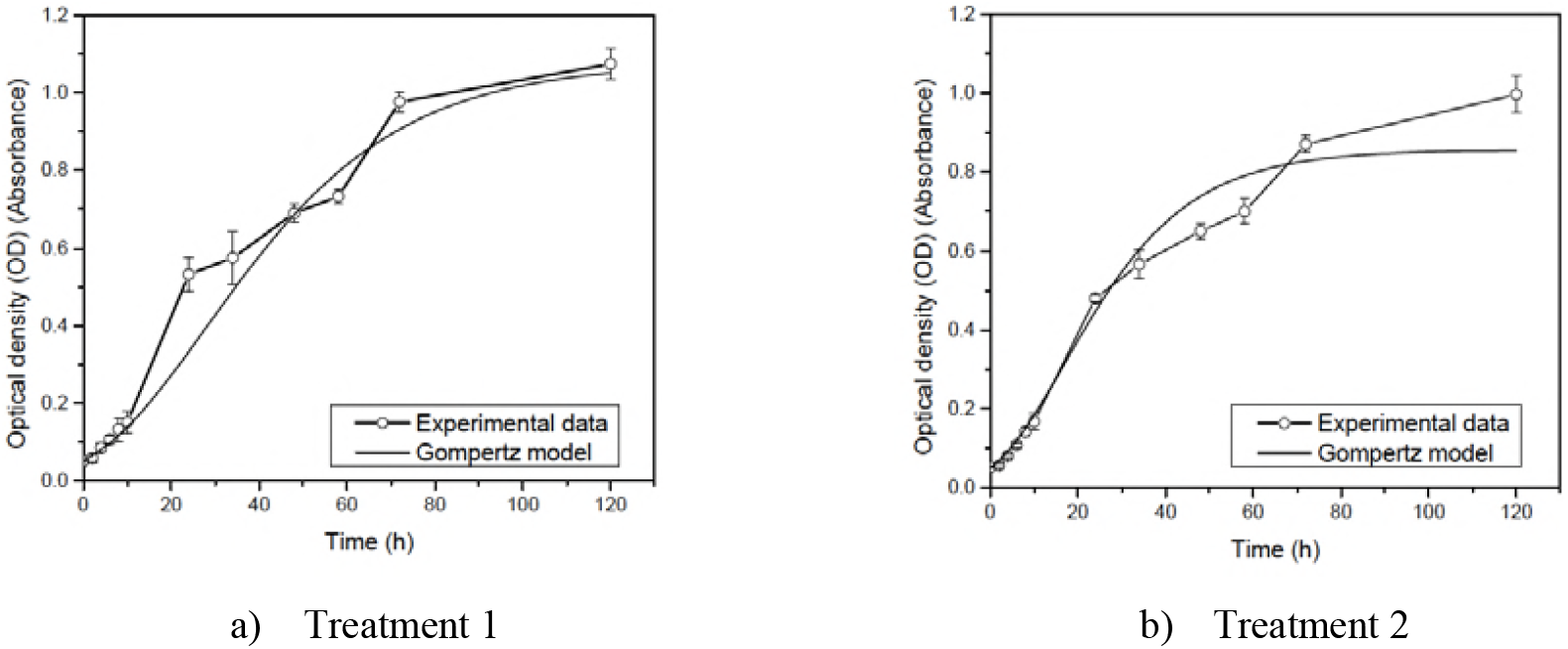
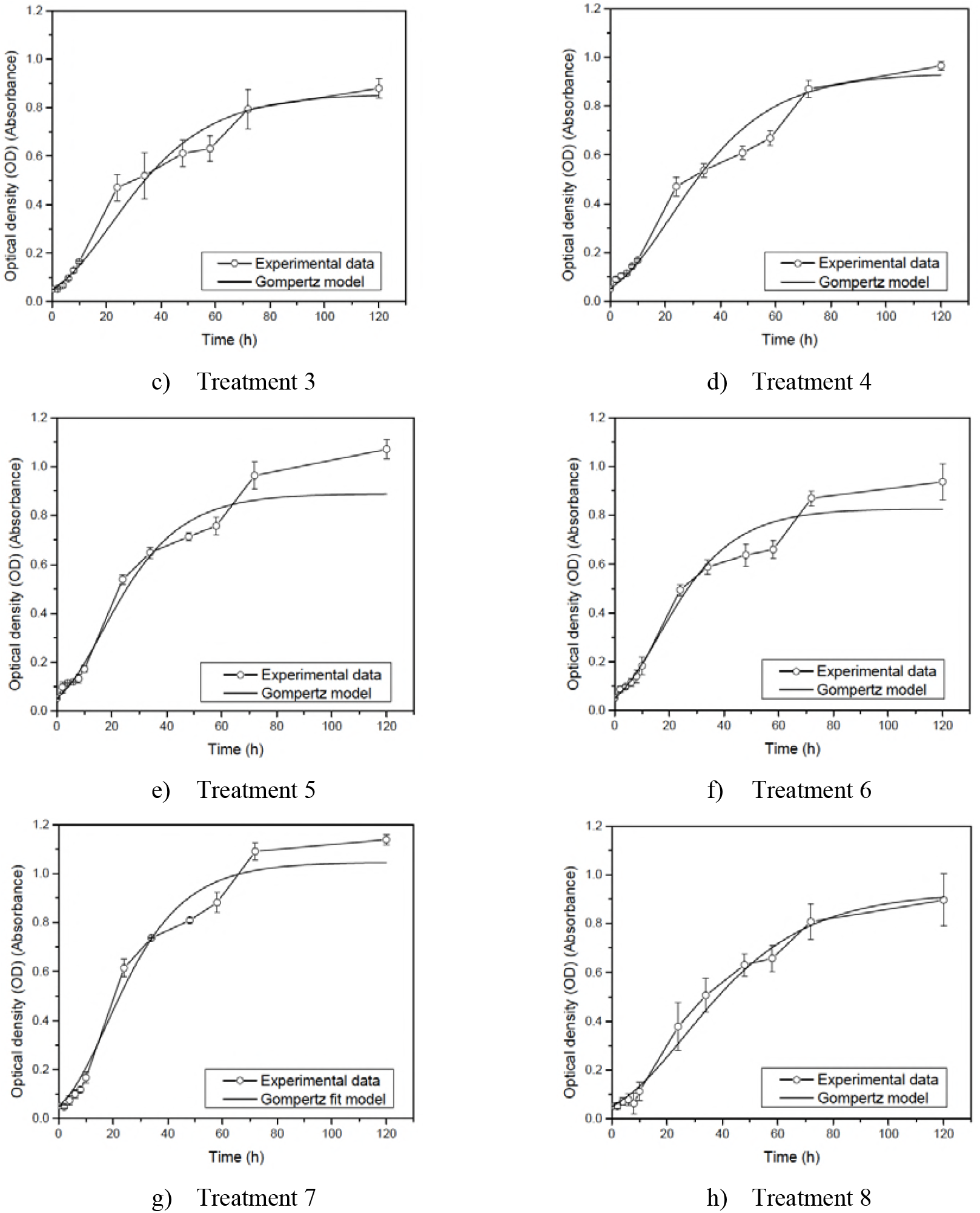
Effect of glucose concentration, initial pH, KH_2_PO_4_ concentration and MgSO_4_ concentration on growth kinetic of *K. medellinensis* NBRC 3288. straight line: modified Gompertz’s model fitted.

**Table 2.** Levels factors evaluated in screening design and results

| Treatment | $X_1$ | $X_2$ | $X_3$ | $X_4$ | Optical density ($\text{OD}_{600}$) | Maximum growth rate ($\mu_m, \text{h}^{-1}$) |
| --- | --- | --- | --- | --- | --- | --- |
| 1 | 2 | 5.0 | 0.05 | 0 | 0.97 | 0.11 |
| 2 | 4 | 5.0 | 0.05 | 0.05 | 0.87 | 0.12 |
| 3 | 4 | 5.0 | 0 | 0 | 0.79 | 0.12 |
| 4 | 2 | 3.5 | 0 | 0 | 0.87 | 0.09 |
| 5 | 2 | 3.5 | 0.05 | 0.05 | 0.96 | 0.10 |
| 6 | 4 | 3.5 | 0.05 | 0 | 0.87 | 0.10 |
| 7 | 2 | 5.0 | 0 | 0.05 | 1.09 | 0.15 |
| 8 | 4 | 3.5 | 0 | 0.05 | 0.80 | 0.09 |

Pereto’s chart (Figure 2) and analysis of variance (Table 3) shows the effects of factors evaluated on biomass production (OD_600_). According to sign of most significant factor in Pareto’s chart, in our case glucose, will apply the steepest ascent method to find a new center of a new experimental region for RSM (Saravanan et al. 2012). The results of steepest ascent method are shown in Figure 3 and the maximum peak occurs at X_1_=0.5 %(wt./v), X_2_=5.38 and X_4_=0.058 %(wt./v) (X_3_ was maintained at 0.025 %(wt./v)), these conditions will be the new center of the new experimental region to obtain a second order model for the optimum of cell density yield.

**Figure 2.**
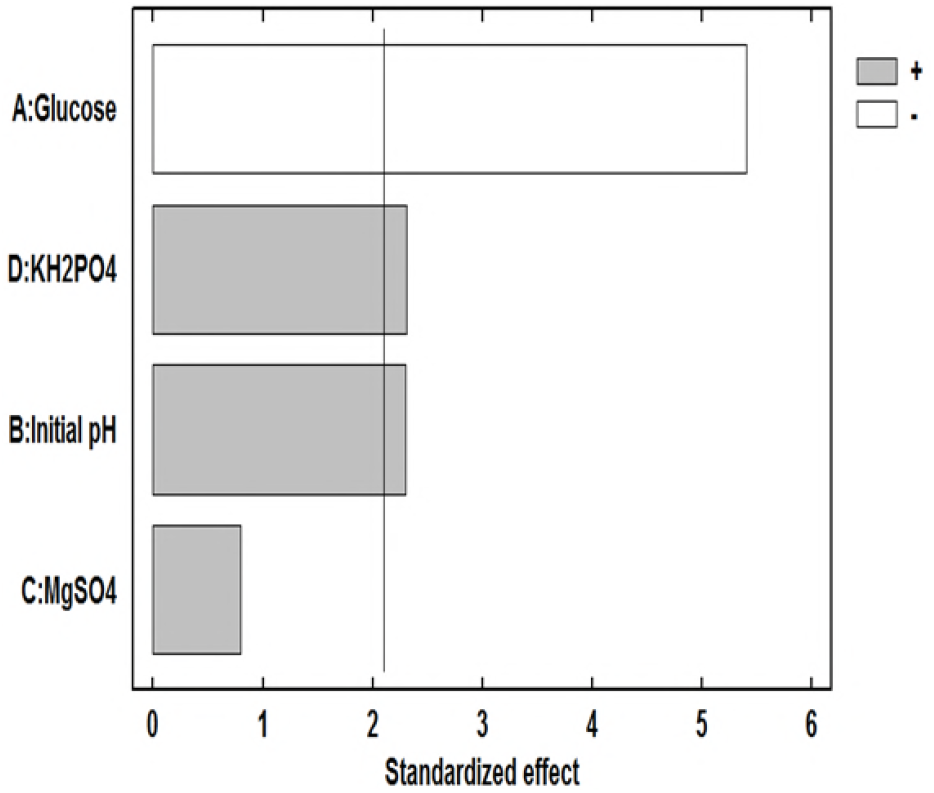
Pareto’s chart where is showed the main effects of factors evaluated. Note that MgSO_4_ was the only not significant factor and the minus sign of glucose means that OD_600_ increases when glucose decrease.

**Figure 3.**
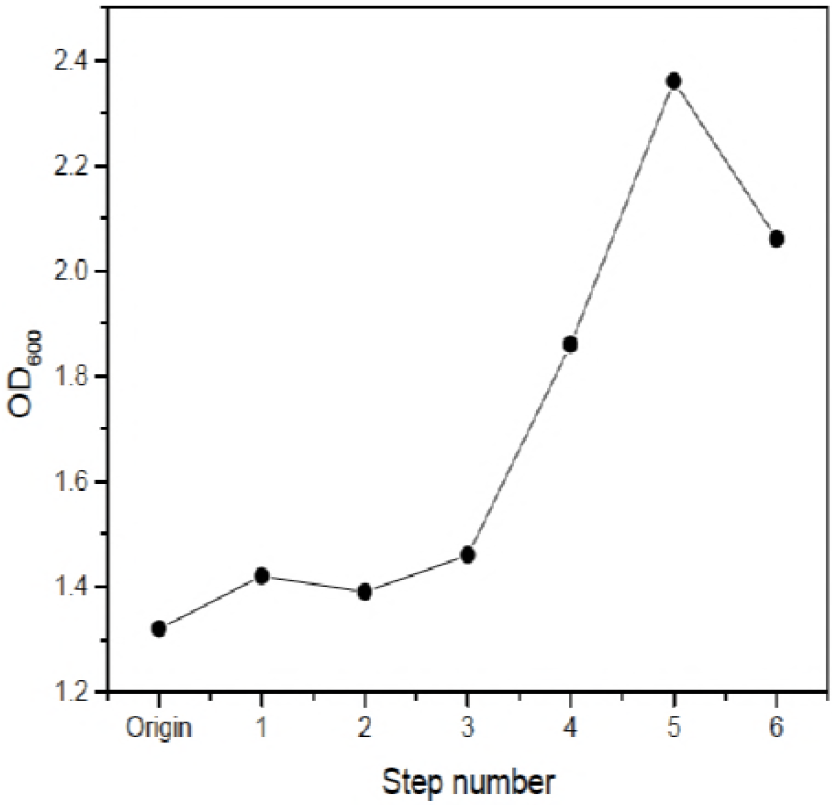
Steepest ascent profile. Note the peak at step 5

**Table 3.**
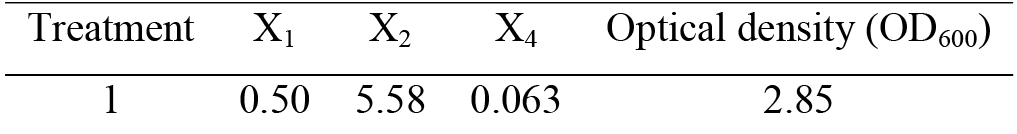
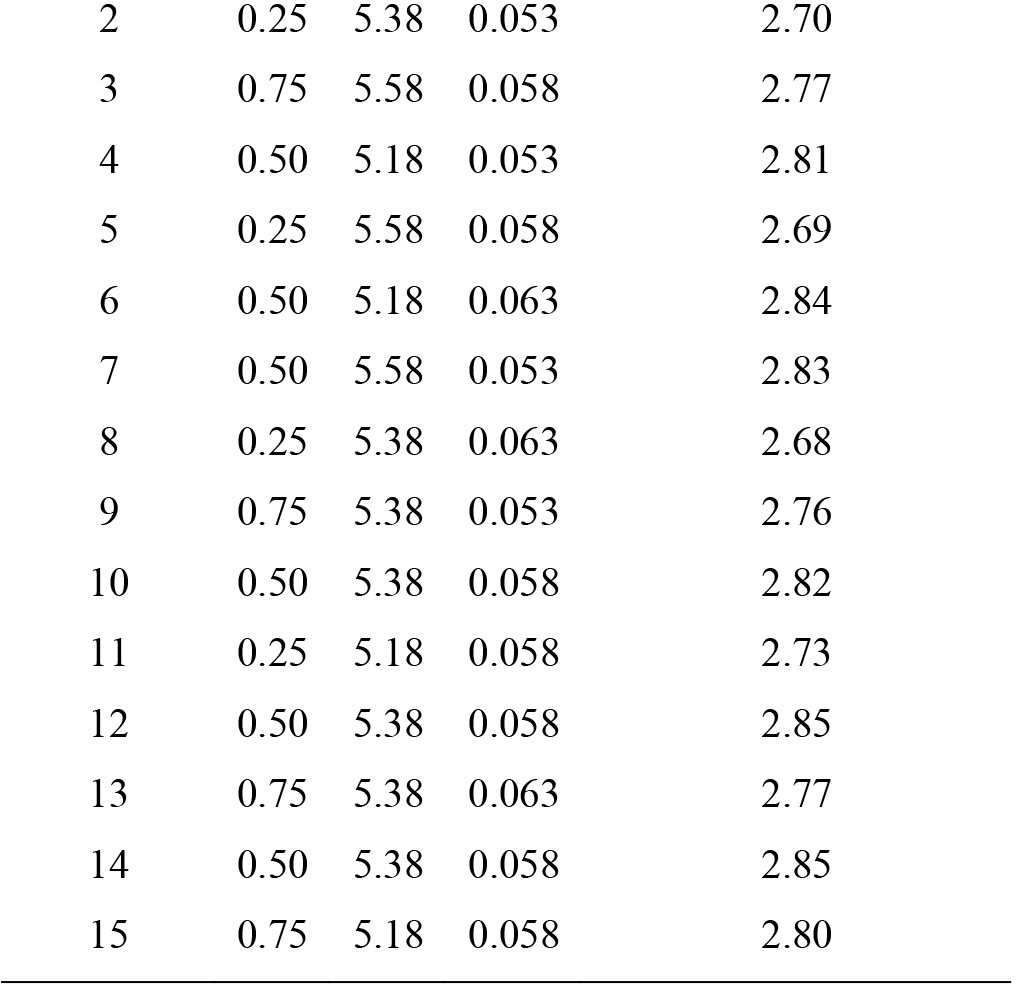
Box-Behnken design and results for cell density yield

### Optimization by response surface methodology

As was mentioned above, a new experimental region will use around the center point glucose (X_1_=0.5 %(wt./v)), initial pH (X_2_=5.38) and KH_2_PO_4_ (X_4_= 0.058 %(wt./v)), selected in fifth step from steepest ascent methodology to optimize biomass production with a Box-Behnken design. The fifteen treatment of Box-Behnken design are in Table 3. Figure 4 and Figure 5 show the response surface plot for biomass production maintained initial pH and KH_2_PO_4_ at one level at time, respectively. At both Figures can be seen that a glucose concentration around 0.5 %(wt./v) was the adequate to optimize biomass production (2.80-2.85 OD600), in fact glucose was the only significant factor to obtain the second order model, as is shown in ANOVA for Box-Behnken design in Table 4. The mathematical second order model is described in equation 2.

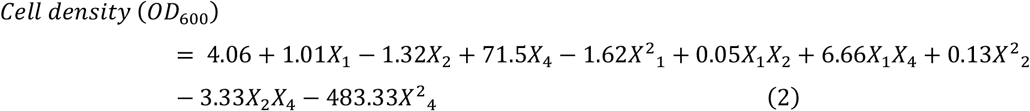

**Figure 4.**
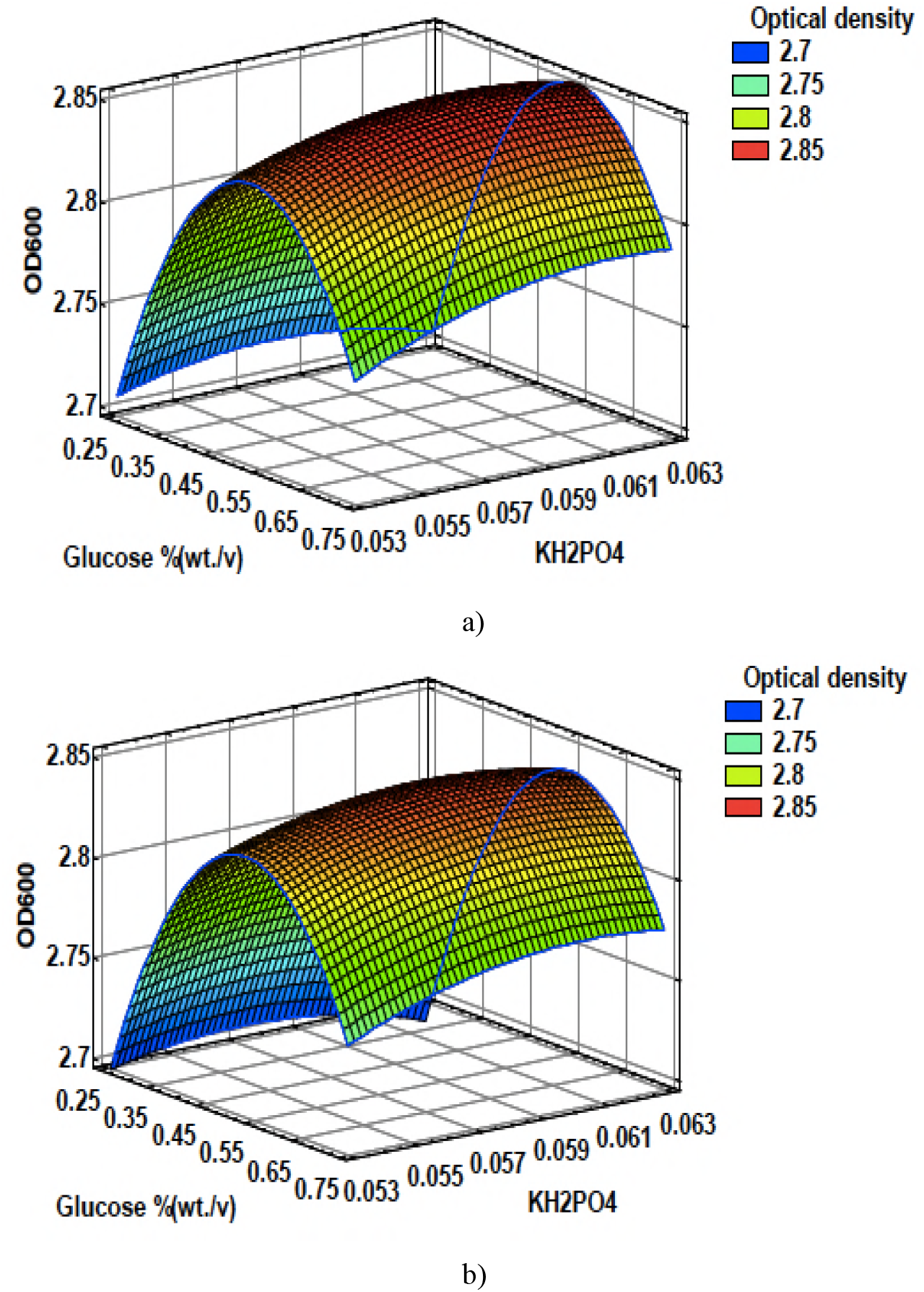
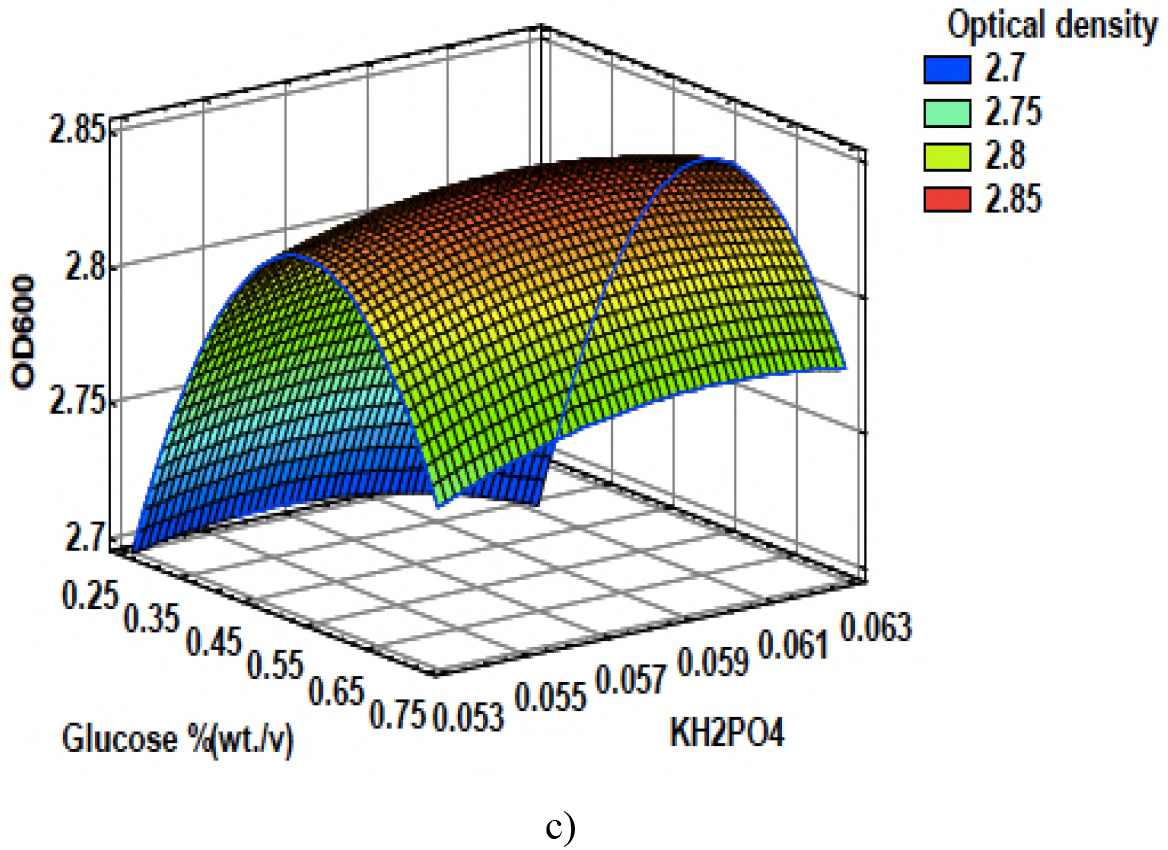
Effect of glucose and KH2 PO4 on cell density yield (OD600). a) pH 5.18, b) pH 5.38 and c) pH 5.58.

**Figure 5.**
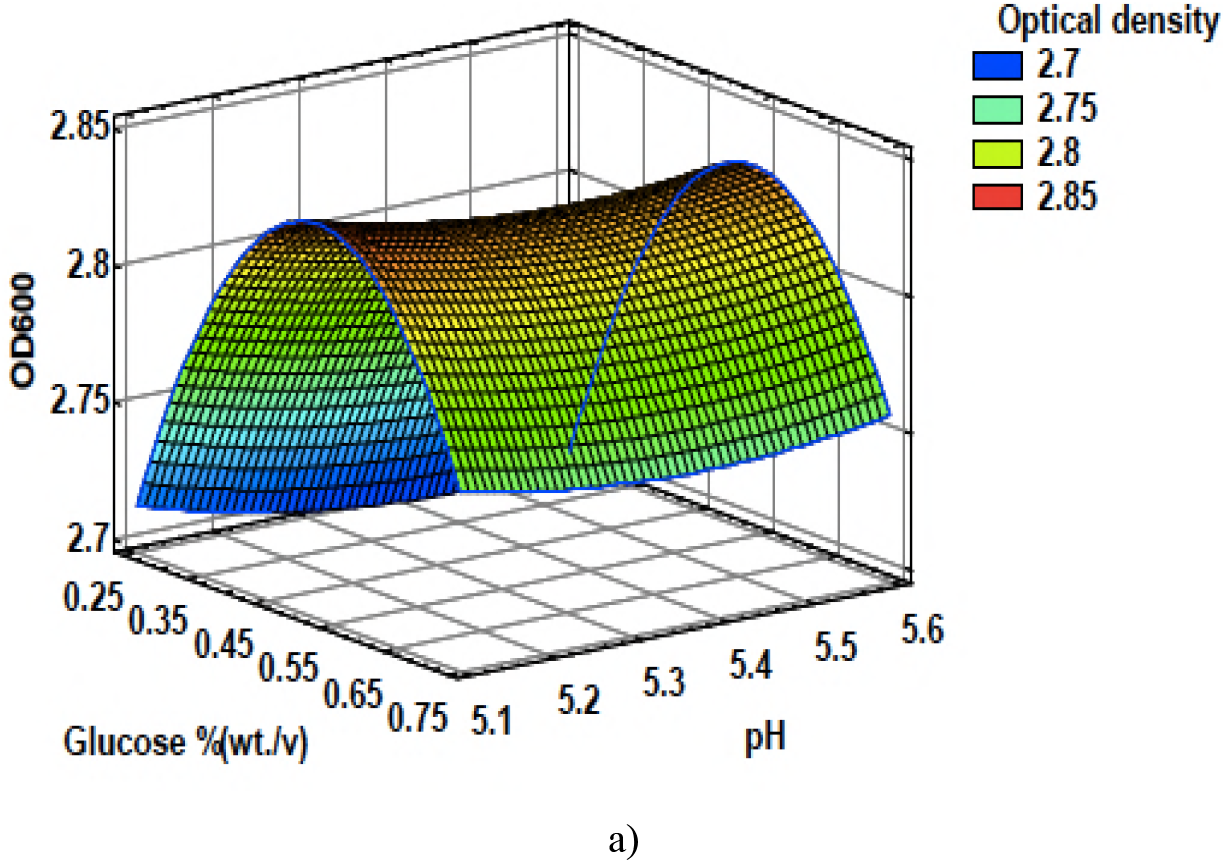
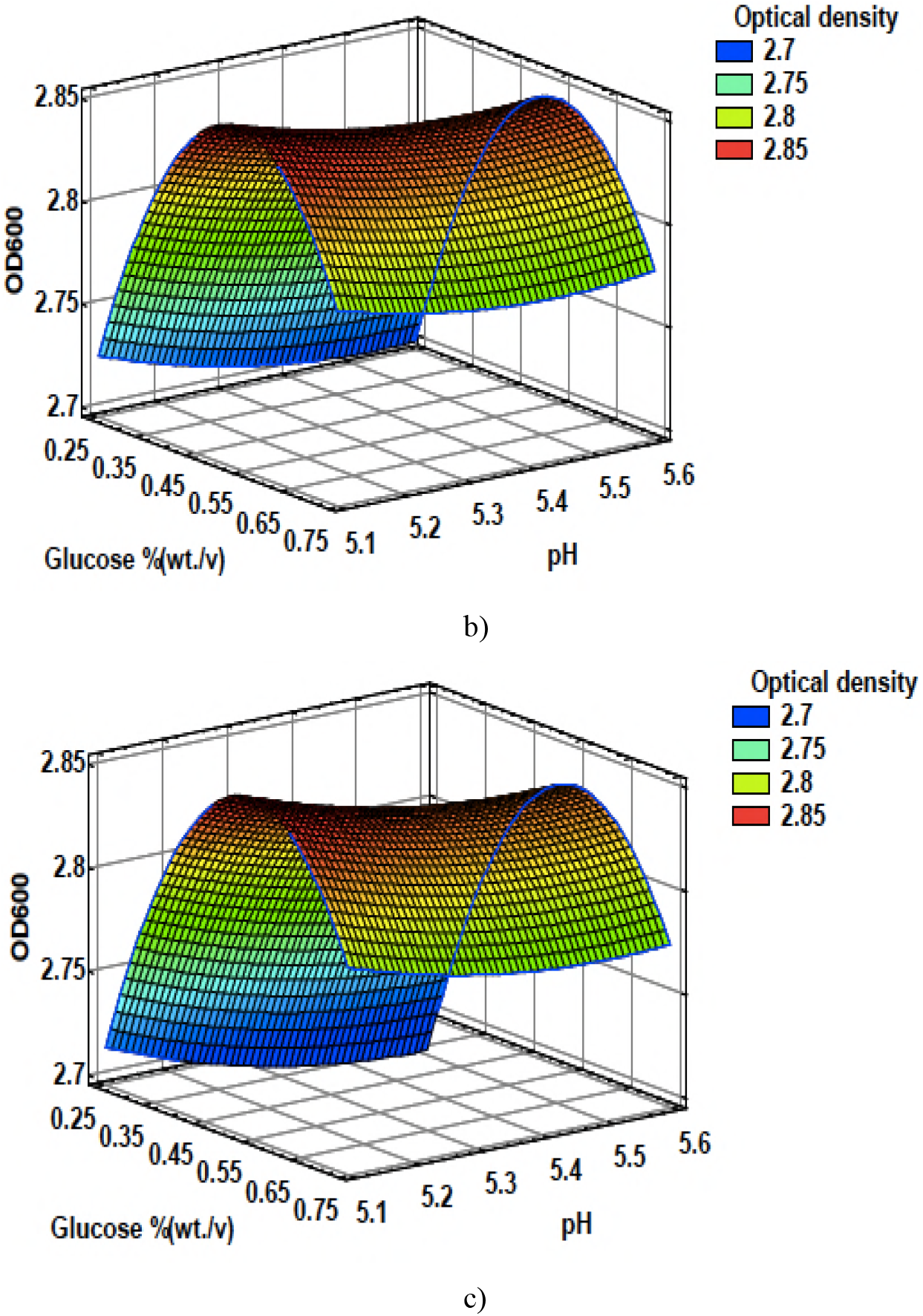
Effect of glucose and initial pH on cell density yield (OD600). a) KH2 PO4 =0.053 %(wt./v), b) KH2 PO4 =0.058%(wt./v) and c) KH2 PO4 =0.063 %(wt./v).

**Table 4.** Analysis of variance of Box-Behnken design for cell density yield of K.

| Source | SS | DF | MS | F-Ratio | P-Value |
| --- | --- | --- | --- | --- | --- |
| X1:Glucose | 0.03 | 1 | 0.03 | 60.39 | 0.00* |
| X2:Initial pH | 0.00 | 1 | 0.00 | 1.80 | 0.22 |
| X4:KH<sub>2</sub>PO<sub>4</sub> | 0.00 | 1 | 0.00 | 0.80 | 0.40 |
| X<sub>2</sub><sub>1</sub> | 0.11 | 1 | 0.11 | 217.45 | 0.00* |
| X1*X2 | 0.00 | 1 | 0.00 | 0.14 | 0.71 |
| X1*X4 | 0.00 | 1 | 0.00 | 1.60 | 0.25 |
| X<sub>2</sub><sub>2</sub> | 0.00 | 1 | 0.00 | 0.62 | 0.46 |
| X2*X4 | 0.00 | 1 | 0.00 | 0.26 | 0.63 |
| X<sub>2</sub><sub>4</sub> | 0.00 | 1 | 0.00 | 3.10 | 0.12 |
| Lack of fit | 0.04 | 27 | 0.00 |  | 0.07 |
| Pure error | 0.00 | 6 | 0.00 |  |  |

SS: Square Sum; DF: Degree of Freedom; MS: Mean Square $R^2=79.07\%$ , $R^2_{adj}=72.09\%$
\*Significant at the 95% confidence level

Finally, a results confirmation was done to verify if model prediction actually was accurate. The Figure 6 shows the growth profile and the the μ_m_ value obtained with the optimum culture medium was 0.23 h^−1^. A faster cell growth was obtained and therefore the stationary phase was reached quickly, value of OD_600_ at 72 h was 2.98.

**Figure 6.**
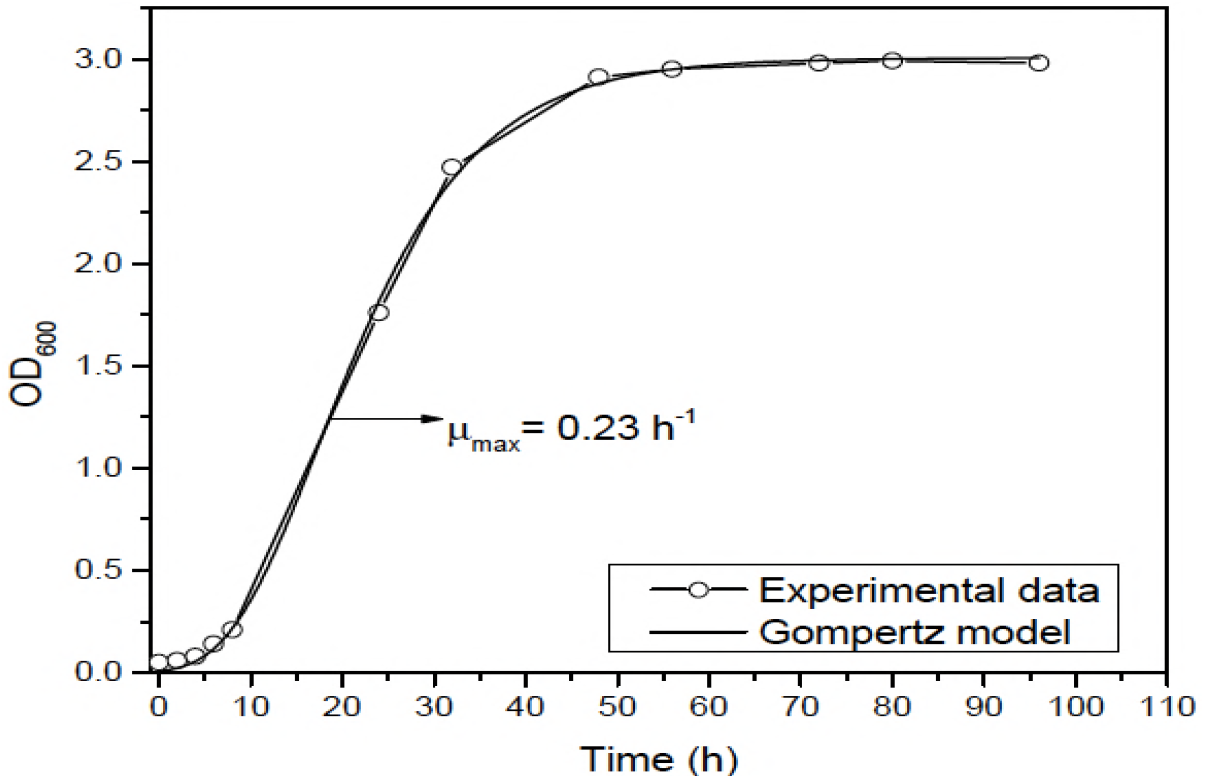
Growth profile at optimum culture medium conditions. Here the growth phases are more notorious and the modified Gompertz model fitted at 99.99 %

## Discussion

In Figure 1 at 72 h starts deceleration of cell growth that is in line to results in static conditions obtained by Molina-Ramírez et al. (Molina-Ramírez et al. 2017) and agitated conditions obtained by Yoshinaga, Tonouchi, Watanabe (Yoshinaga et al. 1997) and Okiyama, Shirae, Kano, Yamanaka (Okiyama et al. 1992) with an strain belong to *Komagataeibacter* genus. These results suggest that fermentation mode (static or agitated) did not affect growth profile, but agitated mode allows obtain a higher cell density yield compared to static mode, in agitated mode a biomass measured by was 1.091(OD_600_) against 0.690 (OD_600_) obtained in static fermentation mode (data not shown). As was mentioned above, 72 h is the age where the *K. medellinensis* NBRC 3288 begins deceleration stage and match with begins of BC production according to Molina-Ramírez et al. (Molina-Ramírez et al. 2017), these results suggest that an inoculum of 72 h, would be the most appropriate to use in the further BC production process (Chao et al. 2000; Bae and Shoda 2004). Likewise, Table 2 shows the results of maximum growth rate (μ_m_, h^−1^) in each treatment evaluated and as it was expected, the highest mm was obtained in the treatment with the highest biomass, which granted that culture medium of treatment 7 is the best to improve cell density yield of *K. medellinensis* NBRC 3288. Results demonstrated that glucose, initial pH and KH2PO4 were significant while MgSO4 was not significant in cell density yield of *K. medellinensis* NBRC 3288 because is known that yeast extract contains traces of Mg^2+^ (Grant and Pramer 1962) and the effect of addition of MgSO_4_ to culture medium is shielded, otherwise KH_2_PO_4_ was significant because is the only source of K+, which is very important to bacterial cells metabolism (Nierhaus 2014). In the case of glucose, in the optimization stage, it was the only significant factor on cell density yield of *K. medellinensis* NBRC 3288 showing a better effect when its concentration was lower. But, cell density yield was not only improved in this work but, that μm was increased from 0.15 h^−1^ until 0.23 h^−1^ (34.78 % of improvement), this because, probably, exponential growth is the sum of overall reactions in the cell and it is controlled by the reaction lead by the enzyme with the highest affinity for glucose which is the only carbon and energy source in the culture medium (Monod 1949; Shehata and Man-1971).

The ANOVA results showed that model explain properly the 72.09 % of data and using optimization function of STATGRAPHICS^®^ was obtained the optimum values of significant factors: glucose, 0.545 %(wt./v); initial pH, 5.18 and KH_2_PO_4_, 0.059 %(wt./v), predicting an optimum value of OD_600_ of 2.85. To verify these results, experiments were performed by triplicated and the results were 2.98 OD_600_, a very near value to predicted by model only 4.26 % above. Due to μm was increased too, this means that *K. medellinensis* NBRC 3288 growth faster in this culture conditions and the inoculum for BC production do not need to wait until 72 h as we have spoken before but until 48 h because at this point start cell growth deceleration allowing reduction in the time that could improve the productivity of the whole process of BC production.

So the composition of the new culture medium is: glucose, 0.54 %(wt./v); peptone, 0.5 %(wt./v); yeast extract, 0.5 %(wt./v); KH_2_PO_4_, 0.059 %(wt./v); MgSO_4_, 0.025 %(wt./v); NaH_2_PO_4_, 0.267 %(wt./v) and initial pH, 5.18 (adjusted with citric acid, 0.2 %(wt./v)) with an improvement of 68.21 % in biomass production of *K. medellinensis* NBRC 3288 measured by OD_600_.

The culture conditions for inoculum achievement, a previous stage of BC production process, was optimized by improvement of cell density of *K. medellinensis* NBRC 3288 and an improvement of μ_m_, making faster this stage of the whole process. Besides a decreasing of cost by reduction of glucose concentration was reached as a plus in this work, allowing money save in futures fermentation.

## Acknowledgement

The authors acknowledge funding from Research Center for Investigation and Development (CIDI) from the Universidad Pontificia Bolivariana.

## Compliance with Ethical Standards

### Funding

This study was funded by the Science, Technology and Innovation Administrative Department of the Colombian Government (COLCIENCIAS) (grant # 672 of 2014).

### Ethical approval

This article does not contain any studies with human participants performed by any of the authors.

## Reference

Bae S, Shoda M (2004) Bacterial Cellulose Production by Fed-Batch Fermentation in Molasses Medium. Biotechnol Prog 20:1366–1371. doi: 10.1021/bp0498490

Bezerra MA, Santelli RE, Oliveira EP, Villar LS, Escaleira LA (2008) Response surface methodology (RSM) as a tool for optimization in analytical chemistry. Talanta 76:965–977. doi: 10.1016/j.talanta.2008.05.019

Castro C, Cleenwerck I, Trcek J, Zuluaga R, De Vos P, Caro G, Aguirre R, Putaux JL, Ganan P (2013) *Gluconacetobacter medellinensis* sp nov., cellulose-and non-cellulose-producing acetic acid bacteria isolated from vinegar. Int J Syst Evol Microbiol 63:1119–1125. doi: 10.1099/ijs.0.043414-0

Castro C, Zuluaga R, Alvarez C, Putaux JL, Caro G, Rojas OJ, Mondragon I, Ganan P (2012) Bacterial cellulose produced by a new acid-resistant strain of *Gluconacetobacter* genus. Carbohydr Polym 89:1033–1037. doi: 10.1016/j.carbpol.2012.03.045

Chao Y, Ishida T Fau-Sugano Y, Sugano Y Fau-Shoda M, Shoda M (2000) Bacterial cellulose production by *Acetobacter xylinum* in a 50-L internal-loop airlift reactor. Biotechnol Bioeng 68:345–358. doi: 10.1002/(SICI)1097-0290(20000505)68:3<345::aid-bit13>3.0.CO;2-m

Deppenmeier U, Ehrenreich A (2009) Physiology of acetic acid bacteria in light of the genome sequence of *Gluconobacter oxydans*. J Mol Microbiol Biotechnol 16:69–80. doi: 10.1159/000142895

Grant CL, Pramer D (1962) Minor element composition of yeast extract. J Bacteriol 84:869–870.

Gutiérrez Pulido H, Vara Salazar R de la, Gutiérrez González P, Téllez Martínez C, Temblador Pérez M del C (2004) Análisis y diseño de experimentos. McGraw-Hill

Haigler CH, Benziman M (1982) Biogenesis of Cellulose I Microfibrils Occurs by Cell-Directed Self-Assembly in Acetobacter xylinum. In: Cellulose and Other Natural Polymer Systems. Springer US, Boston, MA, pp 273–297

Hajati S, Ghaedi M, Yaghoubi S (2015) Local, cheep and nontoxic activated carbon as efficient adsorbent for the simultaneous removal of cadmium ions and malachite green: Optimization by surface response methodology. J Ind Eng Chem 21:760–767. doi: 10.1016/j.jiec.2014.04.009

Hestrin S, Schramm M (1954) Synthesis of cellulose by *Acetobacter xylinum*. 2. Preparation of freeze-dried cells capable of polymerizing glucose to cellulose. Biochem J 58:345–352.

Molina-Ramírez C, Castro M, Osorio M, Torres-Taborda M, Gómez B, Zuluaga R, Gómez C, Gañán P, Rojas O, Castro C (2017) Effect of Different Carbon Sources on Bacterial Nanocellulose Production and Structure Using the Low pH Resistant Strain *Komagataeibacter medellinensis*. Materials (Basel) 10:639. doi: 10.3390/ma10060639

Monod J (1949) The Growth of Bacterial Cultures. Annu Rev Microbiol 3:371–394. doi: 10.1146/annurev.mi.03.100149.002103

Nierhaus KH (2014) Mg2+, K+, and the ribosome. J Bacteriol 196:3817–9. doi: 10.1128/JB.02297-14

Okiyama A, Shirae H, Kano H, Yamanaka S (1992) Bacterial cellulose I. Two-stage fermentation process for cellulose production by *Acetobacter aceti*. Food Hydrocoll 6:471–477. doi: 10.1016/S0268-005X(09)80032-5

Ramos A, Boels IC, de Vos WM, Santos H (2001) Relationship between glycolysis and exopolysaccharide biosynthesis in *Lactococcus lactis*. Appl Environ Microbiol 67:33–41. doi: 10.1128/AEM.67.1.33-41.2001

Ross P, Mayer R, Benziman M (1991) Cellulose biosynthesis and function in bacteria. Microbiol Rev 55:35–58.

Santos SM, Carbajo JM, Gómez N, Quintana E, Ladero M, Sánchez A, Chinga-Carrasco G, Villar JC (2016) Use of bacterial cellulose in degraded paper restoration. Part II: application on real samples. J Mater Sci 51:1553–1561. doi: 10.1007/s10853-015-9477-z

Saravanan P, Muthuvelayudham R, Rajesh Kannan R, Viruthagiri T (2012) Optimization of cellulase production using *Trichoderma reesei* by RSM and comparison with genetic algorithm. Front Chem Sci Eng 6:443–452. doi: 10.1007/s11705-012-1225-1

Shah N, Ul-Islam M, Khattak WA, Park JK (2013) Overview of bacterial cellulose composites: A multipurpose advanced material. Carbohydr Polym 98:1585–1598. doi: http://dx.doi.org/10.1016/j.carbpol.2013.08.018

Shehata TE, Marr AG (1971) Effect of Nutrient Concentration on the Growth of Escherichia coli. J Bacteriol 107:210–216.

Shi F, Xu Z, Cen P (2006) Optimization of γ-polyglutamic acid production by *Bacillus subtilis* ZJU-7 using a surface-response methodology. Biotechnol Bioprocess Eng 11:251–257. doi: 10.1007/BF02932039

Shi Z, Zhang Y, Phillips GO, Yang G (2014) Utilization of bacterial cellulose in food. Food Hydrocoll 35:539–545.

Thomas S, Visakh PM, Mathew AP, Cherian B, Leão A, de Souza S, de Olyveira G, Costa L, Brandão C, Narine S (2013) Bacterial Nanocellulose for Medical Implants. In: Advances in Natural Polymers. Springer Berlin Heidelberg, pp 337–359

Yoshinaga F, Tonouchi N, Watanabe K (1997) Research Progress in Production of Bacterial Cellulose by Aeration and Agitation Culture and Its Application as a New Industrial Material. Biosci Biotechnol Biochem 61:219–224. doi: 10.1271/bbb.61.219

Zeng X, Small DP, Wan W (2011) Statistical optimization of culture conditions for bacterial cellulose production by *Acetobacter xylinum* BPR 2001 from maple syrup. Carbohydr Polym 85:506–513. doi:http://dx.doi.org/10.1016/j.carbpol.2011.02.034

Zhang K (2013) Illustration of the development of bacterial cellulose bundles/ribbons by *Gluconacetobacter xylinus* via atomic force microscopy. Appl Microbiol Biotechnol 97:4353–4359. doi: 10.1007/s00253-013-4752-x

Zwietering MH, Jongenburger I, Rombouts FM, van’t Riet K (1990) Modeling of the bacterial growth curve. Appl Environ Microbiol 56:1875–81.

